# Negative correlation between mean female age at first birth and rates of molecular evolution in primates, with predictions of the reproductive age in early hominins

**DOI:** 10.1101/2020.08.09.232769

**Authors:** Piotr Bajdek

## Abstract

mtDNA-based phylogenetic trees of the order Primates were constructed by the minimum evolution (ME) and maximum likelihood (ML) methods. Branch lengths were compared with the mean female age at first birth in the taxa studied. Higher reproductive age in females triggers a lower number of generations through time and, on average, at the molecular level smaller evolutionary distances between related taxa. However, this relationship is significant when the phylogeny is resolved by the ME method rather than the ML method. Reliability of the minimum evolution approach is discussed. In contrast to most studies, the ME tree recovers *Tarsius bancanus* (Tarsiiformes) as a member of the Strepsirrhini, which phylogeny is supported by a strong branch length–reproductive age relationship and which is proposed as a novel heuristic method to test phylogeny. However, branches of certain taxa on the constructed phylogenetic tree show anomalous lengths relative to the mean female age at first birth, such as e.g. the human branch. As estimated in this paper, early members of the human lineage have likely reproduced at higher rates than modern humans, some forms possibly giving first birth at the mean age of 10–12 years, which is more comparable to the mean age at first birth in extant gorillas than to that typical of living humans and chimpanzees. Probable early reproduction in human ancestors is also supported by the comparably more evolved mitochondrial DNA in Denisovans than in modern humans, and by a smaller body mass in most fossil hominins, which often triggers fast maturation in primates.

## 1. INTRODUCTION

The molecular clock concept constitutes one of the principal tools in phylogenetic studies and allows for construction of time-calibrated trees (Bajdek, 2019). However, the rate of molecular evolution vary among lineages (Wu and Li, 1985; Li and Tanimura, 1987; Yi, 2013; Moorjani et al., 2016) which makes assessing divergence dates problematic (Tsantes and Steiper, 2009). Several causes have been proposed to explain variable molecular rates, such as differences in body size and metabolism leading to a higher rate of DNA damage in small-bodied taxa (Martin and Palumbi, 1993; Fontanillas et al., 2007) as well as changes in DNA replication or repair mechanism (Britten, 1986) and sperm competition intensity (Wong, 2014). Notwithstanding, most researchers postulate that the rate of molecular evolution is influenced primarily by the generation time effect (Wu and Li, 1985; Li and Tanimura, 1987; Elango et al., 2006; Tsantes and Steiper, 2009; Perry et al., 2011).

The rates of molecular evolution in primates and other lifeforms have been studied by applying rather few methods to resolve the phylogeny and branch length distances. The maximum likelihood principle and its derivatives are currently the most frequently used methods (Tsantes and Steiper, 2009; Moorjani et al., 2016), whereas in most of the early studies on the rates of substitution the neighbor-joining algorithm was used (Elango et al., 2006; Kim et al., 2006; Perry et al., 2011). It is remarkable that virtually all recent studies on the phylogeny of primates are based on the maximum likelihood and Bayesian approaches (Chatterjee et al., 2009; Perelman et al., 2011; Finstermeier et al., 2013; Pozzi et al., 2014; Wu et al., 2017), although the maximum parsimony method was also used (Martín-Peciña et al., 2019).

On the contrary to the current trends in phylogenetics, it is arguable that a distance-based principle for resolving phylogeny can be more suitable in studies on the rates of molecular evolution. In order to test this thesis, primate phylogenetic trees resolved by the maximum likelihood and minimum evolution methods are here analyzed by plotting their total path lengths with the mean female age at first birth in the taxa studied. Possible implications for the phylogeny of primates and in particular the evolutionary history hominines are also discussed.

## 2. MATERIAL AND METHODS

### 2.1. Study of the reproductive age in primates

The mean age at reproduction might best correlate with branch lengths on the phylogenetic tree but reliable estimates thereof are unavailable for most primate taxa. Instead, the mean age at first birth for various primate species was retrieved from literature, similarly as done by Tsantes and Steiper (2009). During the selection of taxa for analyses it was necessary to balance between evenly sampling different parts of the primate phylogenetic tree, and including species which reproductive biology is well-studied and corresponding mtDNA sequences are available in the NCBI database. Whenever possible, values correspond to animals studied in the wild. 25 living primate species were selected in total (Table 1).

**Table 1.**
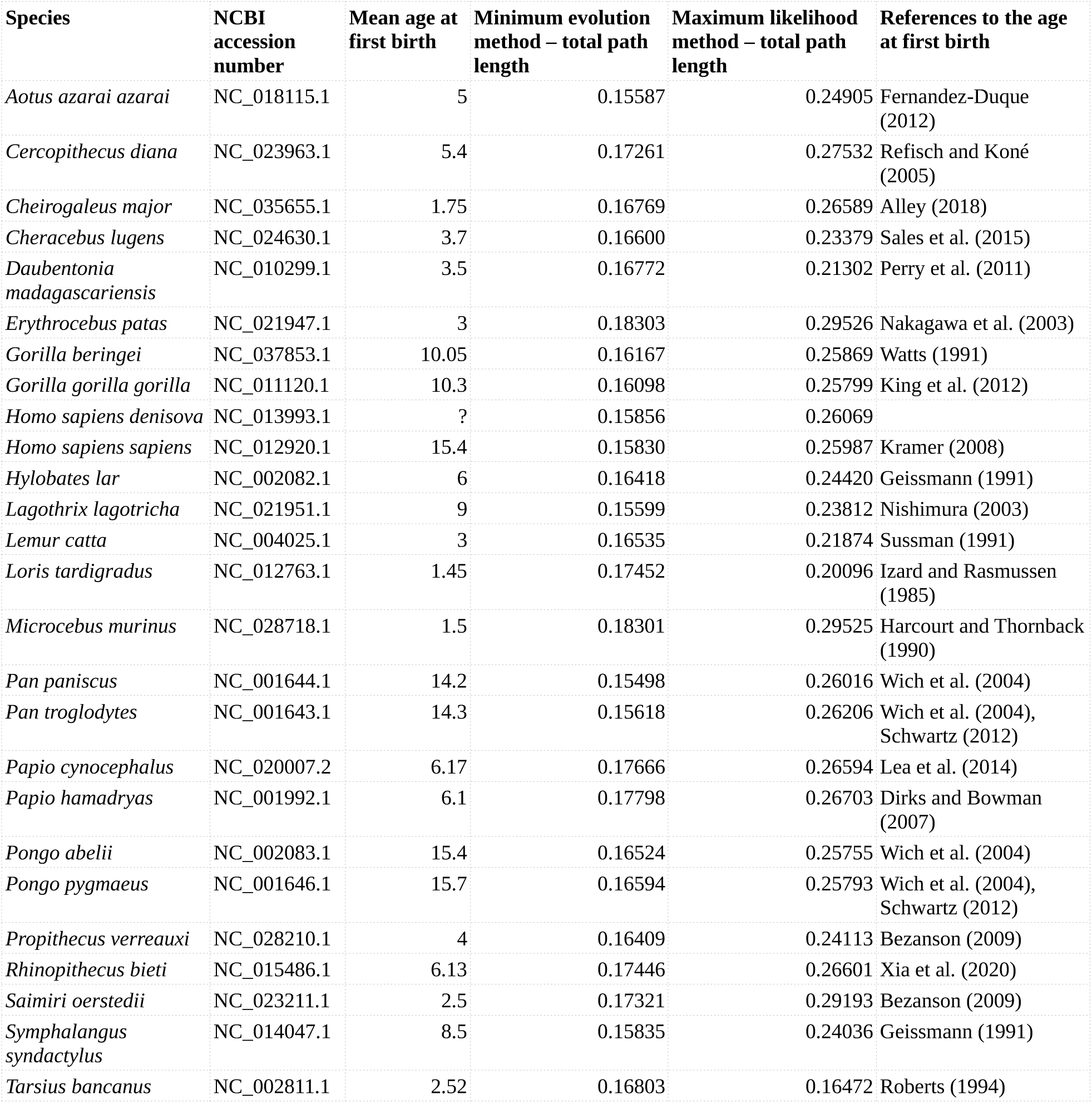
Data analyzed in this study and calculated total path lengths.

| Species | NCBI accession number | Mean age at first birth | Minimum evolution method – total path length | Maximum likelihood method – total path length | References to the age at first birth |
| --- | --- | --- | --- | --- | --- |
| *Aotus azarai azarai* | NC_018115.1 | 5 | 0.15587 | 0.24905 | Fernandez-Duque (2012) |
| *Cercopithecus diana* | NC_023963.1 | 5.4 | 0.17261 | 0.27532 | Refisch and Koné (2005) |
| *Cheirogaleus major* | NC_035655.1 | 1.75 | 0.16769 | 0.26589 | Alley (2018) |
| *Cheracebus lugens* | NC_024630.1 | 3.7 | 0.16600 | 0.23379 | Sales et al. (2015) |
| *Daubentonia madagascariensis* | NC_010299.1 | 3.5 | 0.16772 | 0.21302 | Perry et al. (2011) |
| *Erythrocebus patas* | NC_021947.1 | 3 | 0.18303 | 0.29526 | Nakagawa et al. (2003) |
| *Gorilla beringei* | NC_037853.1 | 10.05 | 0.16167 | 0.25869 | Watts (1991) |
| *Gorilla gorilla gorilla* | NC_011120.1 | 10.3 | 0.16098 | 0.25799 | King et al. (2012) |
| *Homo sapiens denisova* | NC_013993.1 | ? | 0.15856 | 0.26069 |  |
| *Homo sapiens sapiens* | NC_012920.1 | 15.4 | 0.15830 | 0.25987 | Kramer (2008) |
| *Hylobates lar* | NC_002082.1 | 6 | 0.16418 | 0.24420 | Geissmann (1991) |
| *Lagothrix lagotricha* | NC_021951.1 | 9 | 0.15599 | 0.23812 | Nishimura (2003) |
| *Lemur catta* | NC_004025.1 | 3 | 0.16535 | 0.21874 | Sussman (1991) |
| *Loris tardigradus* | NC_012763.1 | 1.45 | 0.17452 | 0.20096 | Izard and Rasmussen (1985) |
| *Microcebus murinus* | NC_028718.1 | 1.5 | 0.18301 | 0.29525 | Harcourt and Thornback (1990) |
| *Pan paniscus* | NC_001644.1 | 14.2 | 0.15498 | 0.26016 | Wich et al. (2004) |
| *Pan troglodytes* | NC_001643.1 | 14.3 | 0.15618 | 0.26206 | Wich et al. (2004), Schwartz (2012) |
| *Papio cynocephalus* | NC_020007.2 | 6.17 | 0.17666 | 0.26594 | Lea et al. (2014) |
| *Papio hamadryas* | NC_001992.1 | 6.1 | 0.17798 | 0.26703 | Dirks and Bowman (2007) |
| *Pongo abelii* | NC_002083.1 | 15.4 | 0.16524 | 0.25755 | Wich et al. (2004) |
| *Pongo pygmaeus* | NC_001646.1 | 15.7 | 0.16594 | 0.25793 | Wich et al. (2004), Schwartz (2012) |
| *Propithecus verreauxi* | NC_028210.1 | 4 | 0.16409 | 0.24113 | Bezanson (2009) |
| *Rhinopithecus bieti* | NC_015486.1 | 6.13 | 0.17446 | 0.26601 | Xia et al. (2020) |
| *Saimiri oerstedii* | NC_023211.1 | 2.5 | 0.17321 | 0.29193 | Bezanson (2009) |
| *Symphalangus syndactylus* | NC_014047.1 | 8.5 | 0.15835 | 0.24036 | Geissmann (1991) |
| *Tarsius bancanus* | NC_002811.1 | 2.52 | 0.16803 | 0.16472 | Roberts (1994) |

The mean female age of 15.4 years at first birth used as representative of modern humans (*Homo sapiens sapiens*) corresponds to the Pumé/Yaruro tribe of primarily hunter-gatheres from the Venezuelan Orinoquia, whose reproduction was studied by Kramer (2008). Although the mean female age at first birth in present-day industrialized nations is notably higher (Mathews et al., 2002), it is often assumed that nonagricultural human populations are more suitable for human–other primates biological comparisons (see Alberts et al., 2013; Willems and van Schaik, 2017).

### 2.2. Involvement of fossil taxa

Despite the reproductive age in extinct taxa is unknown, one mtDNA sequence belonging to Denisovans was involved in the analyses in aim to study more in detail the human branch. Neanderthal mtDNA was excluded from the analyses due to its high similarity to that of *Homo sapiens sapiens*. The study of subtle differences in branch lengths may require a reasonable evolutionary distance between taxa in order to obtain a reliable signal. *Homo heidelbergensis* was also excluded, whose available mtDNA sequence is slightly incomplete. Lastly, to counter the node density effect which can distort the branch lengths (Venditti et al., 2006; Hugall and Lee, 2007), it was important to balance the phylogenetic tree by even sampling (e.g., two human species vs. two chimpanzee species vs. two gorilla species).

### 2.3. Phylogenetic analyses

Evolutionary analyses involved 26 mtDNA sequences of primates including 1 fossil and 25 extant taxa (Table 1), which were downloaded from the National Center for Biotechnology Information, U.S. National Library of Medicine. The analyses were conducted in MEGA X version 10.1.8 (Kumar et al., 2018) on Fedora 32.

Sequence alignment was performed by the MUSCLE algorithm (Edgar, 2004). Constructing the phylogenetic trees all positions containing gaps and missing data were eliminated (complete deletion option) and there was a total of 15095 positions in the final datasets. Two distinct methods were used:

a. The Minimum Evolution (ME) method (Rzhetsky and Nei, 1992). The Neighbor-joining algorithm (Saitou and Nei, 1987) was used to generate the initial tree. The evolutionary distances were computed using the Maximum Composite Likelihood method (Tamura et al., 2004) and are in the units of the number of base substitutions per site. The ME tree was searched using the Close-Neighbor-Interchange (CNI) algorithm (Nei and Kumar, 2000) at a search level of 1. The optimal tree with the sum of branch length = 2.26173061 is shown (Fig. 1A; Fig. 3). The phylogeny was tested by 10000 bootstrap replicates (Felsenstein, 1985).
b. The Maximum Likelihood (ML) method and Tamura-Nei model (Tamura and Nei, 1993). Initial trees for the heuristic search were obtained automatically by applying Neighbor-Join (Saitou and Nei, 1987) and BioNJ (Gascuel, 1997) algorithms to a matrix of pairwise distances estimated using the Tamura-Nei model, and then selecting the topology with superior log likelihood value. The tree with the highest log likelihood (−176040.91) is shown (Fig. 1B). The phylogeny was tested by 1000 bootstrap replicates (Felsenstein, 1985).

## 3. RESULTS AND DISCUSSION

### 3.1. Comparison of the ME and ML methods

The phylogenetic trees of the order Primates constructed by the use of the minimum evolution (ME) and maximum likelihood (ML) methods are compared in Fig. 1. The topologies of the trees are mostly the same, although the phylogenies of the Lemuriformes and Platyrrhini were resolved differently. The tree constructed via the ME approach has higher bootstrap test results (Fig. 1).

**Fig. 1.**
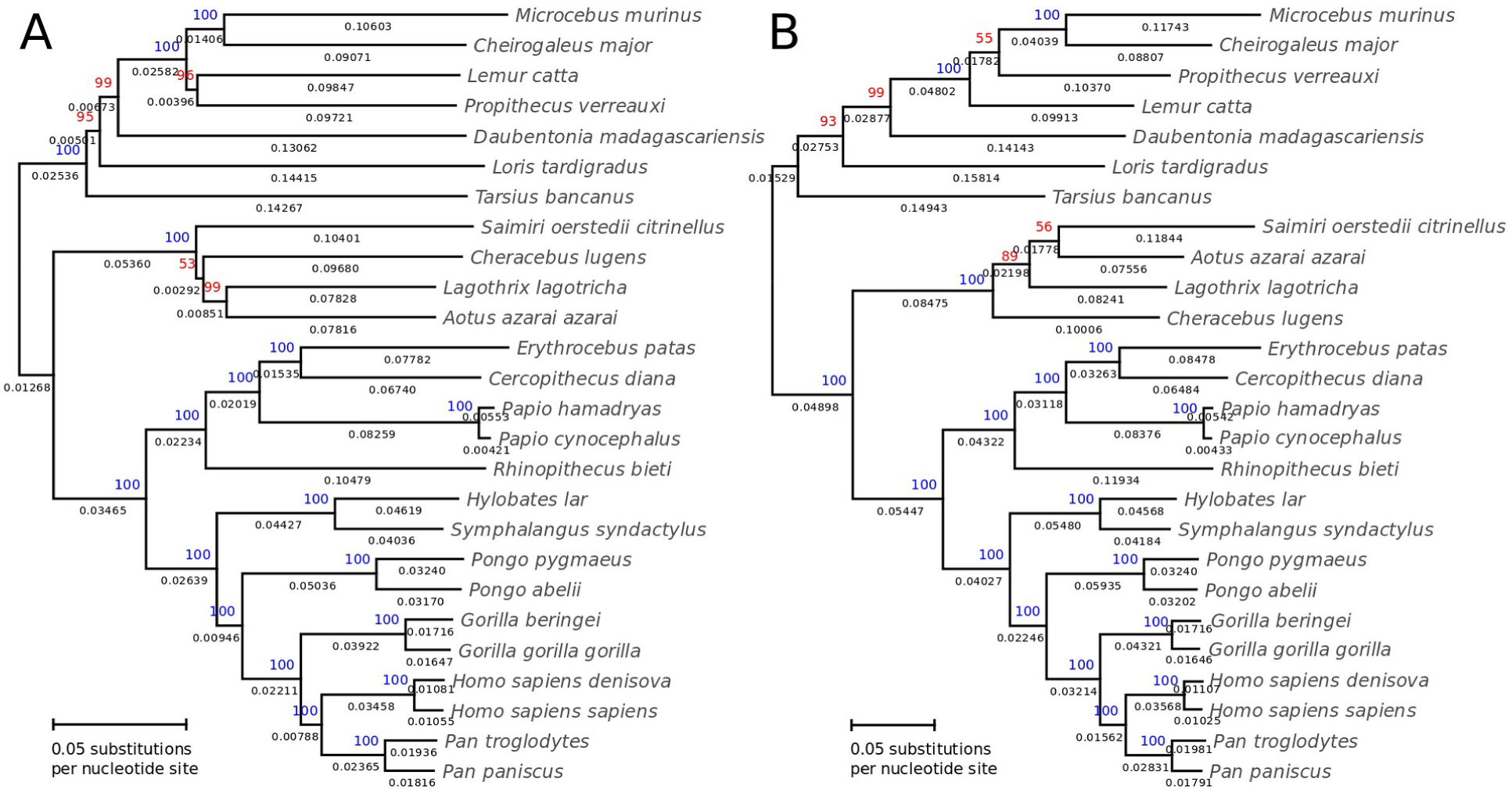
Minimum evolution (A) and maximum likelihood (B) mtDNA-based phylogenetic trees of primates. The percentage of replicate trees in which the associated taxa clustered together in the bootstrap test are shown in blue (when = 100%) and red (when < 100%). Branch lengths are shown below the branches and are in the units of the number of base substitutions per site.

**Fig. 2.**
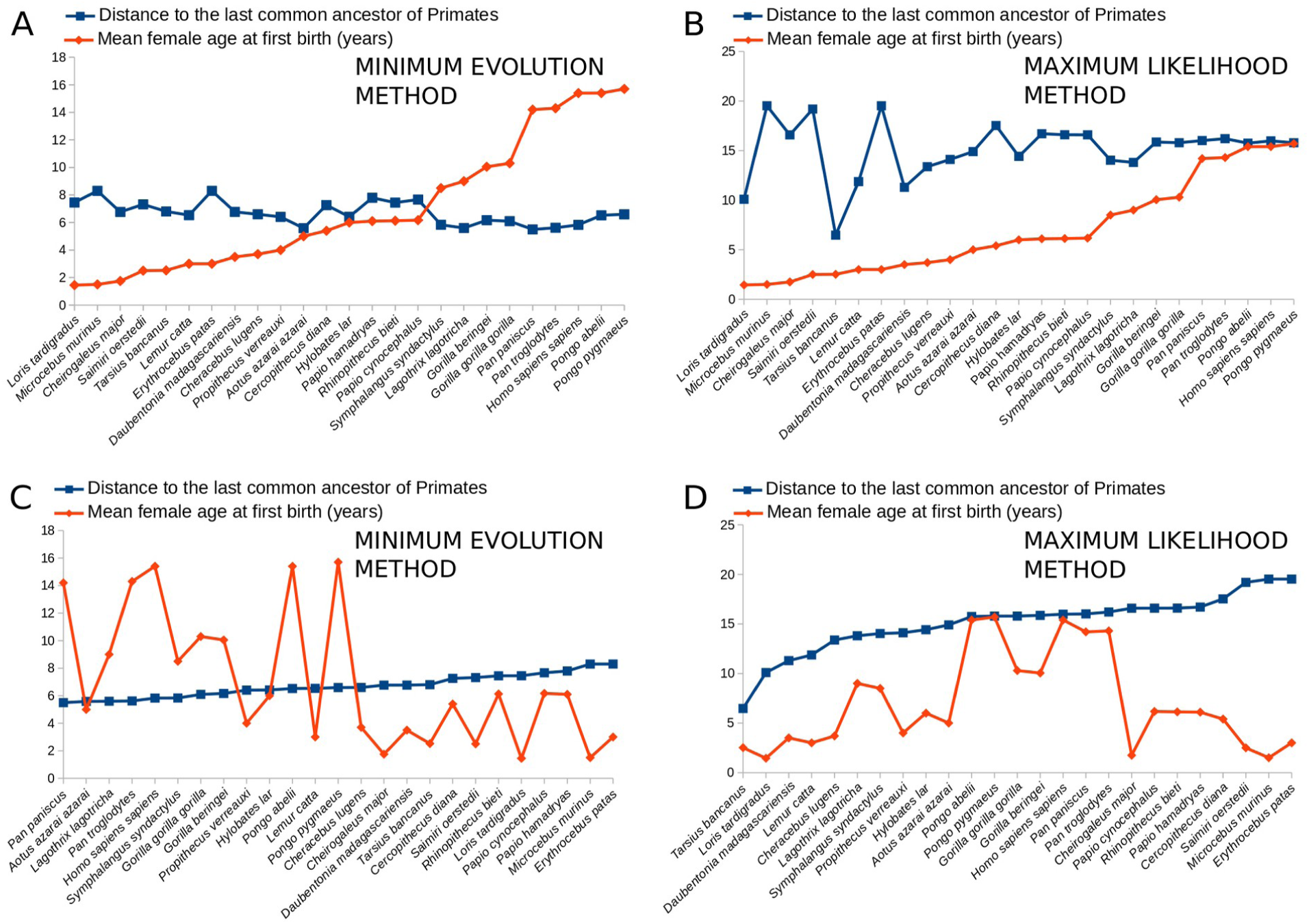
Relationship between the mean female age at first birth (red line) and the total path length (blue line) based on the minimum evolution (A, C) and maximum likelihood (B, D) methods. For illustrative purposes the evolutionary distances are algorithmic calculated by the formula: *total path length * 100 – 10*, based on the Table 1. Note that whereas the ages at first reproduction and evolutionary distances are negatively correlated based on the ME method (Pearson’s r = -0,582; Spearman’s ρ = -0,589), this correlation is missing when the ML method is applied (Pearson’s r = 0,178; Spearman’s ρ = 0,059).

**Fig. 3.**
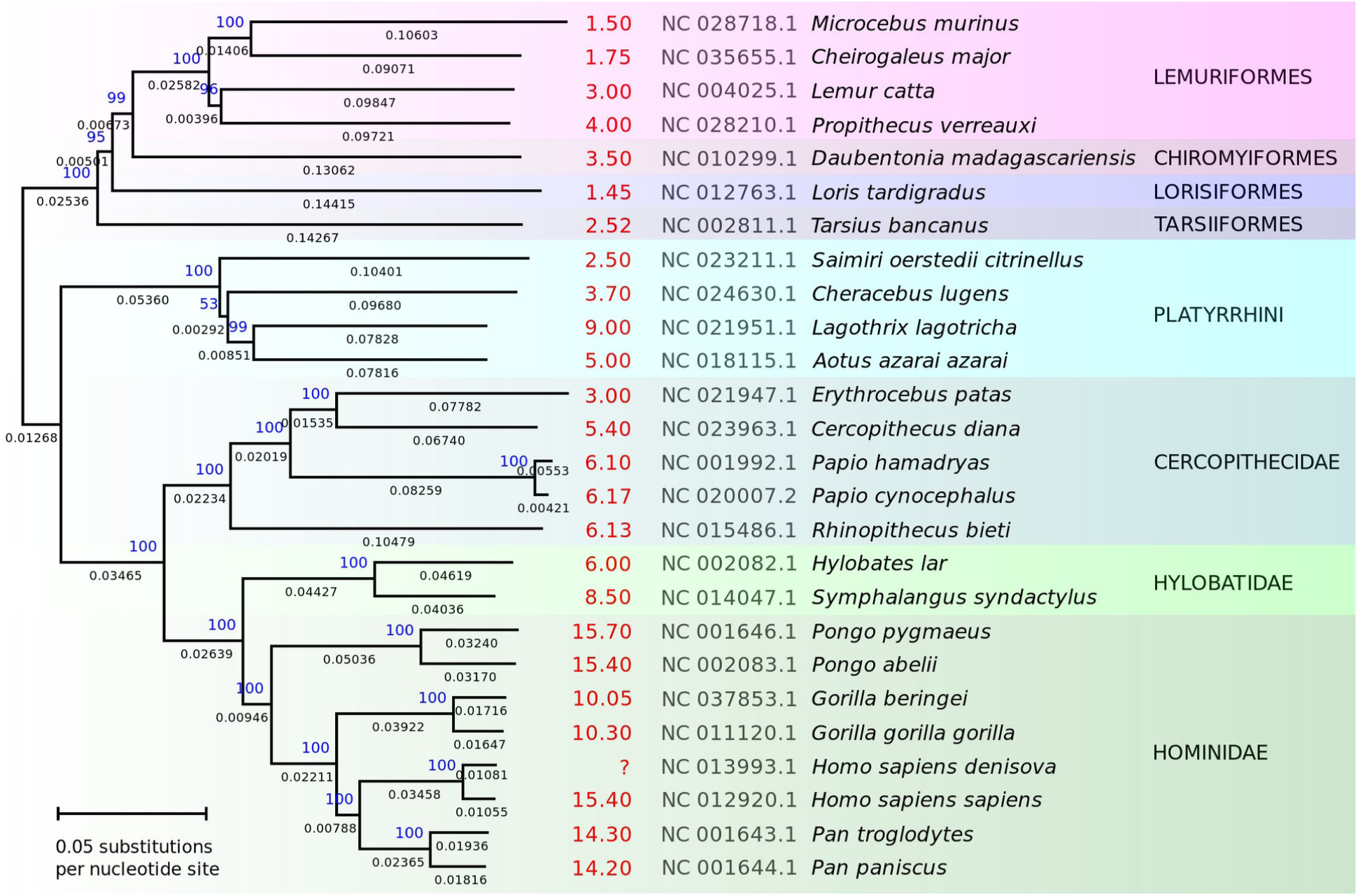
Minimum evolution tree of the order Primates. Note that the mean female age at first birth (in red; years) is negatively correlated with the total path lengths. NCBI accession numbers are provided in gray and bootstrap test results in blue.

Total path lengths to the last common ancestor of crown-group primates were calculated for both trees (Table 1) and plotted with the mean female age at first birth (Fig. 2). When the phylogeny is resolved by the ME method, the reproductive age negatively correlates with the distances: Pearson’s r = -0,582, Spearman’s ρ = -0,589 (Fig. 2A, C). On the contrary, there is no clear correlation when the ML method is applied: Pearson’s r = 0,178, Spearman’s ρ = 0,059 (Fig. 2B, D). Calculated correlation coefficients are provided in Table 2.

**Table 2.** Correlation coefficients between the distances to the last common ancestor of the order Primates and the mean female age at first birth. Note the moderate–high negative correlation when the phylogeny is resolved by the ME method and the general lack of this relationship when the ML method is used.

| Minimum Evolution (ME) |  |  |  |  |
| --- | --- | --- | --- | --- |
| | Pearson's r | Spearman's $\rho$ | Linear regression for<br>y = age at first birth | Linear regression for<br>x = branch length |
| Primates | -0,582 | -0,589 | $y = -335,522 * x + 62,975$ | $x = -0,001 * y + 0,174$ |
| Strepsirrhini | -0,737 | -0,786 | $y = -113,362 * x + 21,81$ | $x = -0,005 * y + 0,182$ |
| Haplorhini | -0,575 | -0,473 | $y = -299,586 * x + 58,339$ | $x = -0,001 * y + 0,175$ |
| *G. beringei* + *G. gorilla* +<br>*P. paniscus* + *P. troglodytes* | -0,986 | -0,8 | $y = -690,077 * x + 121,557$ | $x = -0,001 * y + 0,176$ |
| Maximum Likelihood (ML) |  |  |  |  |
| | Pearson's r | Spearman's $\rho$ | Linear regression for<br>y = age at first birth | Linear regression for<br>x = branch length |
| Primates | 0,178 | 0,059 | $y = 29,072 * x + -0,321$ | $x = 0,001 * y + 0,244$ |
| Strepsirrhini | -0,261 | -0,036 | $y = -6,137 * x + 3,934$ | $x = -0,011 * y + 0,257$ |
| Haplorhini | -0,245 | -0,253 | $y = -68,215 * x + 26,455$ | $x = -0,001 * y + 0,268$ |
| *G. beringei* + *G. gorilla* +<br>*P. paniscus* + *P. troglodytes* | 0,888 | 0,8 | $y = 1161,716 * x + -289,514$ | $x = 0,001 * y + 0,251$ |

The minimum evolution (ME) method is based on the assumption that the tree with the smallest sum of branch length estimates is most likely to be the true one (Rzhetsky and Nei, 1992, 1993). First, the neighbor-joining (NJ) tree is obtained by the Saitou and Nei’s (1987) method and then the tree with the minimum value of the total sum of branch lengths is searched by examining all trees that are closely related to the NJ tree (Rzhetsky and Nei, 1992, 1993). Results of the experiment in this study corroborate this method as an effective approach for assessing the “true” branch lengths. Higher mean age at first birth triggers smaller evolutionary distances between related taxa, as most easily explained by a lower number of generations, but this relationship is better visible through the ME than the ML method (Fig. 2; Table 2).

### 3.2. Phylogenetic position of the Tarsiiformes

The phylogenetic position of tarsiers (Tarsiiformes) is contentious (Yoder, 2003) since molecular studies support their placement as a sister group to either the suborder Strepsirrhini (e.g., Murphy et al., 2001; Waddell et al., 2001; Eizirik et al., 2004; Chatterjee et al., 2009; Huang, 2012) or the Haplorhini (e.g., Ziętkiewicz et al., 1999; Perelman et al., 2011; Springer et al., 2012; Finstermeier et al., 2013; Hartig et al., 2013; Pozzi et al., 2014). Their placement within the Haplorhini appears to be the currently prevailing viewpoint (see López-Torres, 2018) and is supported by the lack of the ability to synthesise the vitamin C in *Tarsius bancanus* which is shared with simians (Pollock and Mullin, 1987), and also other putative haplorhine synapomorphies (Yoder, 2003).

On the other hand, mtDNA analyses in this study recover Tarsiiformes as a sister group to Strepsirrhini by the use of both the ME and ML methods (Fig. 1). It should be noted that *Tarsius bancanus* is more deeply placed on the strepsirrhine branch by the ME method (Fig. 1A), whereas the primate phylogeny in most of the mtDNA-based previous studies was resolved by applying the ML and Bayesian algorithms (Finstermeier et al., 2013; Pozzi et al., 2014). Notwithstanding, Chatterjee et al. (2009) recovered *Tarsius* within the suborder Strepsirrhini on the basis of mitochondrial and nuclear genes and Bayesian phylogenetic methods.

Most importantly, when the ME approach is applied, the total path length of *Tarsius bancanus* appears to correlate with members of the Strepsirrhini (Fig. 3). Correlation coefficient between the total path lengths and the mean female age at first birth for Strepsirrhini, including *T. bancanus*, is as high as Pearson’s r = -0,737 and Spearman’s ρ = -0,786 (Table 2). For example, the total path on *T. bancanus* is slightly shorter than the paths of *Microcebus murinus* and *Loris tardigradus* which give first birth earlier, and slightly longer than the total paths of *Propithecus verreauxi, Lemur catta* and *Daubentonia madagascariensis* which on average give first birth at a higher age (Fig. 3).

In contrast, placing *Tarsius bancanus* as a sister taxon to the suborder Haplorhini requires its branch to be more than 0.01268 substitutions per site or 8.89% longer than it is, if this total path length–reproductive age relationship is kept. Arguably, the most parsimonious approach for the study of phylogeny is to seek a scenario requiring the lowest possible amount of evolutionary changes.

### 3.3. The human lineage

#### 3.3.1. Comparison with gorillas and chimpanzees

The present study corroborates the hominoid slowdown hypothesis (Goodman, 1985; Li and Tanimura, 1987; Steiper et al., 2004) as the paths of the families Hominidae and Hylobatidae are fairly short relative to other groups of the Primates, which is more evident resolving the phylogeny by the ME method (Fig. 3). However, in contrast to early results (Li and Tanimura, 1987; Elango et al., 2006) and in accordance with recent studies (Perelman et al., 2011; Finstermeier et al., 2013; Pozzi et al., 2014), the path of *Homo* is strikingly long compared with the paths of *Gorilla* and *Pan*, and the concomitant reproductive age in living humans (Fig. 3). *G. beringei, G. gorilla, P. troglodytes*, and *P. paniscus* are characterized by a high negative correlation coefficient between their total path lengths and the mean female age at first birth: Pearson’s r = -0,986; Spearman’s ρ = -0,8 (Table 2; ME approach). Based on these four species this linear regression equation can be formulated: *y = -690,077 * x + 121,557*, where *y* = mean female age at first birth and *x* = the total path length. If *x* = 0.15830, being the total path length of *H. s. sapiens* (Table 1), then *y* = 12.3178109.

Hence, compared with the rates of molecular evolution in *Gorilla* and *Pan*, the mtDNA of *H. s. sapiens* diverged from that of their last common ancestor at a high rate as if human females on average had given first birth at the age of ∼12.32 years. This is a strikingly lower value than the ∼15.4 years at first parturition found in extant *H. s. sapiens* and also the age of ∼14.2–14.3 in *Pan* (Table 1). Therefore, it appears that certain members of the human lineage have reproduced on average much earlier than modern humans.

Assuming that ∼12.32 years represent the mean value between the first birth age at such earlier stages of the human evolution, and the first birth age of ∼15.4 years in living humans, the former can be estimated as ∼9.24 years: *12.32–(15.4–12.32)=9.24*. This calculation is somewhat simplistic because the shift in reproductive age in human ancestors was almost certainly nonlinear (see chapter 3.3.3.). Nevertheless, this value approximates to the mean age at first birth in gorillas which is estimated as ∼10.05 years for *G. beringei* (see Watts, 1991) and ∼10.3 years for *G. gorilla* (see King et al., 2012). It cannot be ruled out that early members of the human lineage have reproduced at a rate more comparable to that encountered in extant gorillas rather than to that typical of living humans or chimpanzees.

#### 3.3.2. Devisovan mtDNA

The Denisova hominin (Krause et al., 2010) has been variously termed as *Homo denisova* (e.g., Waddell et al., 2011; Gunbin et al., 2012), *Homo altaiensis* (e.g., Zubova et al., 2017), *Homo sapiens denisova* (e.g., Gunbin et al., 2015; Dorado et al., 2018), and *Homo sapiens altaiensis* (e.g., Derevianko, 2011). However, molecular studies reveal that Denisovans interbred with both *Homo sapiens sapiens* (see Reich et al., 2010) and *Homo sapiens neanderthalensis* (see Slon et al., 2018) proving that from the biological viewpoint all these forms belonged to the same species, and should be considered as subspecies of *Homo sapiens* (Linnaeus, 1758). In this paper, the Denisova hominin is termed as *Homo sapiens denisova*.

Early reproduction in human ancestors is supported by the comparably more evolved mtDNA in *H. s. denisova* than in *H. s. sapiens*, which is confirmed by both the ME and ML methods (Fig. 1, Table 1). The mtDNA of *H. s. denisova* is 0.00026 substitutions per site more diverged from that of its last ancestor shared with *H. s. sapiens* than it is the mtDNA of living humans (ME method). Thus, despite the analyzed mtDNA of Denisovans comes from a 30.000– 50.000-years-old fossil (Krause et al., 2010), it appears that there have been more generations in the lineage of *H. s. denisova* than in *H. s. sapiens*, which points out that some fossil hominins might have reproduced at a higher rate than living humans.

The path of *G. beringei* is 0.00549 substitutions per nucleotide site longer than that of *P. troglodytes*, which corresponds to the age difference of 4.25 years at first birth (Fig. 3) and also a ∼10-million-years-long period of separation of lineages (Pozzi et al., 2014). On the other hand, the lineages of *H. s. sapiens* and *H. s. denisova* were estimated to had diverged only ∼1.39 Ma by Pozzi et al. (2014) assuming that *Sahelanthropus tchadensis* (Brunet et al., 2002) was an early member of the human lineage (see Brunet et al., 2005; Parins-Fukuchi et al., 2019). If *0.00549/10Ma=4.25* years, and *0.00026/1.39Ma=x* years, then *x=∼1.45* years of age difference. This suggests that *H. s. denisova* on average have given first birth ∼1.45 years earlier than *H. s. sapiens*, i.e. at the age of ∼13.95. Noteworthy, the earliest births recorded for modern humans are at the age of around 14 years, although such early pregnancy is often associated with adverse biological consequences (Garn et al., 1986; Kramer, 2008).

Tentatively, reproductive age in earlier hominin forms could be also predicted. Assuming that the last common ancestor of *H. s. denisova* and *H. s. sapiens* 1.39 Ma have reproduced at a comparable rate giving first birth at the age of ∼13.95 years, and also a linear shift in reproductive age through time, by extrapolation the earliest forms of *Australopithecus anamensis* (Leakey et al., 1995) living ∼4.15 Ma (White et al., 2006) would have given first birth at the age of ∼11.07: *15.4–(4.15x(1.45/1.39))=∼11.07*. By further extrapolation, the reproductive age typical of *Gorilla* (10.05–10.30 years at first birth; Table 1) would be reached ∼5 Ma, shortly after the final genetic split of the human and chimpanzee lineages understood as the end of hybridization (see Patterson et al., 2006). This is in accordance with the estimation in the chapter above based on the branch length dinstances in *H. s. sapiens, Pan*, and *Gorilla*. Nevertheless, these extrapolations should be treated as an approximation due to the fact that the evolutionary body mass change in the human lineage was nonlinear (see chapter 3.3.3.).

#### 3.3.3. Body size in hominins

Body mass in the human lineage increased through time (Grabowski et al., 2015; Will and Stock, 2015; Jungers et al., 2016), whereas in mammals it tends to be positively correlated with the age of maturity. For example, the siamang (*Symphalangus syndactylus*) is twice the size of other gibbons (Kappelman, 1996) and, on average, gives first birth at a higher age (Geissmann, 1991). Accordingly, the siamang’s (*Symphalangus syndactylus*) path is notably shorter than that of the lar gibbon (*Hylobates lar*), as supported by both the ME and ML methods (Figs 1, 3). Also, the average female body mass in *P. troglodytes* is ∼10 kg lower (∼47.4 kg) than it is in *H. s. sapiens* (57.2 kg) (Kappelman, 1996) and accordingly *P. troglodytes* gives first birth ∼1.1 years earlier than *H. s. sapiens* (Fig. 3; Table 1).

Mean female body mass and mean age at first birth in *H. s. sapiens* (57.2 kg, 15.4 years) and *P. troglodytes* (47.4 kg, 14.3 years) can be tentatively extrapolated on fossil hominins by this regression equation: *y = 0,112 * x + 8,98*, where *y* = age at first birth (years), and *x* = body mass (kg). Grabowski et al. (2015) estimated the mean female body mass for several species of fossil hominins. For example, female *H. erectus* weighted ∼46.3 kg, which suggests the age at first birth = 14.1656 and, interestingly, matches the value of ∼13.95 years estimated herein for *H. s. denisova* based on its mtDNA. Furthermore, the body mass of ∼27.3 kg in female *H. habilis* (Leakey et al., 1964) corresponds to the mean age of ∼12.04 years at first birth.

This all suggests that the relatively small-bodied members of the human lineage have probably reached sexual maturity faster than living humans, which confirms inferences from the branch lengths. Overall, female australopiths weighted around a half of the body mass of female modern humans (Grabowski et al., 2015). However, the evolutionary change in body size in hominins was nonlinear (Jungers et al., 2016). For example, *Homo habilis* was apparently smaller than the some older in geologic age *Australopithecus afarensis* (see Jungers et al., 2016), although their estimated body mass difference is more significant in males than females (Grabowski et al., 2015). Therefore, the evolutionary change in maturity age in the human lineage was almost certainly nonlinear as well.

Irrespectively, gorillas give first birth 4–5 years earlier than humans and chimpanzees but have a much higher body mass. Also, gorillas differ from humans and chimpanzees in having a strong body mass sexual dimorphism (Kappelman, 1996; McHenry, 1996). It has been widely proposed that early members of the human lineage were characterized by such a strong sexual dimorphism and also that this would point out a gorilla-like social and reproductive behavior (Leutenegger and Shell, 1987; Kappelman, 1996; McHenry, 1996). In particular, strong body mass and canine size sexual dimorphism has been suggested for *Australopithecus afarensis* (see Leutenegger and Shell, 1987; Richmond and Jungers, 1995; Plavcan et al., 2005; Gordon et al., 2008; Grabowski et al., 2015), although a few researchers disagreed with these interpretations (Reno et al., 2003, 2010).

## 4. CONCLUSIONS

In spite of the growing popularity of sophisticated approaches for resolving phylogeny, complex methods are not necessarily superior to simple ones. The minimum evolution method has solid theoretical foundations grounded in the Ockham’s razor as it assumes that the most probable phylogeny is that one with the smallest sum of branch lengths. The present study demonstrates a significantly higher negative correlation between the total path lengths and the mean female age at first birth when the mtDNA-based phylogeny of the Primates is resolved by the ME method than the ML method. This calls attention to the ME approach as potentially useful in the studies on the rates of molecular evolution.

Furthermore, assuming that the negative correlation between the total path lengths and the mean age at first birth is not casual but caused by the generation time effect, it is possible that under some circumstances such relationship or lack thereof can be used as a heuristic test of phylogeny. Tree showing a high negative correlation coefficient between these two values is more likely to be the true one than an alternative tree missing it.

Lastly, the study of branch length anomalies allows making evolutionary inferences. For example, branch length distances among living and fossil hominins studied herein suggest that the rate of reproduction in the human lineage has been on average much higher than it is modern humans. It appears that certain members of the human lineage have given first birth at age more comparable to that encountered in gorillas than to that typical of living humans and chimpanzees.

